# Computationally Guided Design of Two Novel Soluble Epoxide Hydrolase Inhibitors

**DOI:** 10.1101/2022.09.13.507703

**Authors:** Elena C. Dere, Simon SK Chu, Teresa Ortega, Peishan Huang, Justin B. Siegel

## Abstract

The enzyme soluble epoxide hydrolase (sEH) has been found to play a role in many ailments such as inflammation, pain, renal function, pulmonary function, hypertension, and diabetes. Multiple sEH inhibitors have been developed to reduce the adverse effects of the ailments. Due to high inhibitory concentrations, there is urgent need for developing improved sEH inhibitors. In this study, two novel sEH inhibitors were designed via computational bioisosteric replacement and chemical intuition with the goal of increasing binding affinity, which can potentially decrease inhibitory concentration. The new drug candidates were found to have improved binding properties compared to existing drugs.

## INTRODUCTION

Soluble epoxide hydrolase (sEH) is an enzyme that is found widely in mammalian tissues, including the brain, liver, heart, spleen, lungs, and kidneys.^1^ They are the primary catalyst for converting epoxyeicosatrienoic acids (EETs) into dihydroxyeicosatetrienoic acids (DHETs) through the conversion of epoxides to diols via hydrolysis.^1^ These EETs are important because they are responsible for vasodilation, vascular smooth muscle cell anti-migratory actions and anti-inflammation among other functions within the body.^2^ The metabolism of EETs has been shown to have adverse effects on inflammation, pain, pulmonary function, hypertension and diabetes.

The use of sEH inhibitors to increase EET levels within the body has significant potential applications for addressing ailments such as inflammation, pain, Alzheimer’s disease, pulmonary fibrosis, inflammatory bowel disease, hypertension, diabetes and more.^3^ Studies have shown that sEH inhibitors have advantages over the more common anti-inflammatory drugs including having a higher potency than most nonsteroidal anti-inflammatory drugs (NSAIDs) and reducing some of the adverse effects of cyclo-oxygenase-2 (COX-2) inhibitors.^2^ These advantages of sEH inhibitors over the typical NSAIDs make them a target of interest in discovering more efficacious anti-inflammatory drugs. Although sEH inhibitors have not yet been approved for use in humans, 1-(1-Propionylpiperidin-4-yl)-3-(4- (trifluoromethoxy)phenyl)urea (TPPU), 1-(1-acetypiperidin-4-yl)-3-adamantanylurea (APAU), and 12-(((tricyclo(3.3.1.13,7)dec-1-ylamino)carbonyl)amino)-dodecanoic acid (AUDA) are three examples of the sEH inhibitors that have been tested in preclinical studies (Figure 1). These studies, conducted on mice and cynomolgus monkeys, indicate that despite being efficacious a relatively high dose was required to elicit sEH inhibition.^4,5,6^ Potency of a drug is highly influenced by the binding affinity, which is the ability of the drug to bind to the target molecule.^7^ In order to better inhibit sEH in humans, it is important to design new drug candidates that have higher binding affinities.

**Figure 1.**
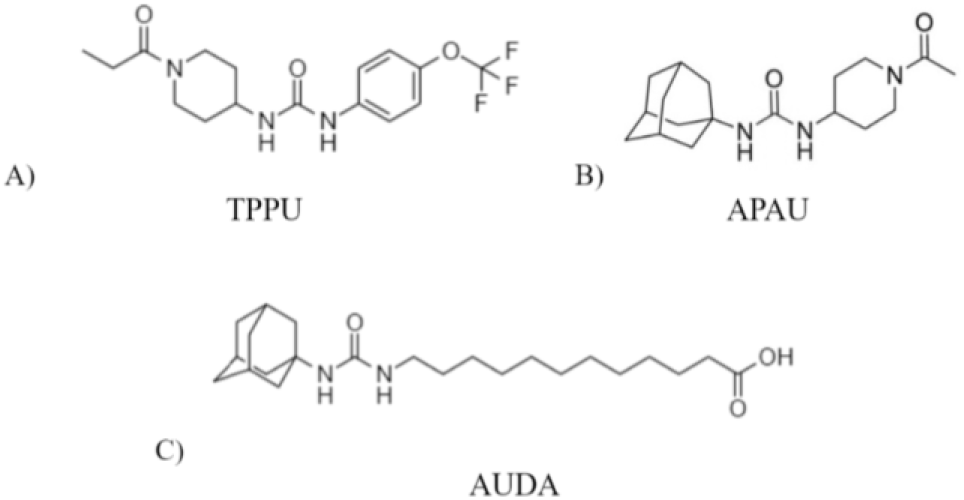
Two-dimensional structures of sEH inhibitors. A) TPPU. B) APAU. C) AUDA.

The primary pharmacophore of these sEH inhibitors has been identified as a derivative of urea, as the urea mimics the epoxide substrate and the transition state of epoxide hydrolysis reaction.^9^ Further studies hypothesize that the pharmacophore (Figure 2) contains an adamantyl structure, a piperidine ring and a polar functional group, such as an ester, ketone, or alcohol, which serves as the secondary pharmacophore.^10^ The adamantyl group improves the pharmacokinetics of the drugs when examining drug concentration within the blood during preclinical trials.^11^ Drugs containing a piperidine structure are found to increase drug absorption and solubility with less plasma protein binding, which enhances its drug-availability to free tissues.^4^ The secondary pharmacophore is linked via a non-polar linker, which is found to increase drug potency.

**Figure 2.**
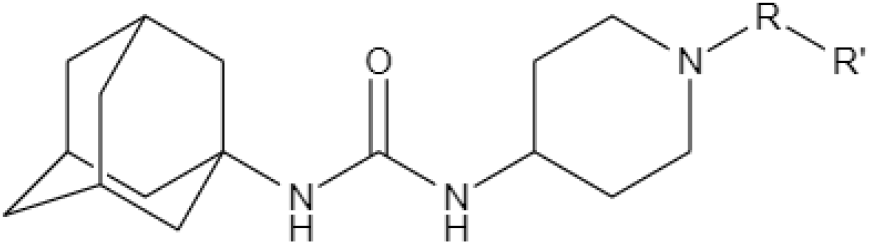
The hypothesized pharmacophore for sEH inhibitors.^12^ The substituents R and R’ represent a non-polar linker and the secondary pharmacophore, respectively.

**Figure 3.**
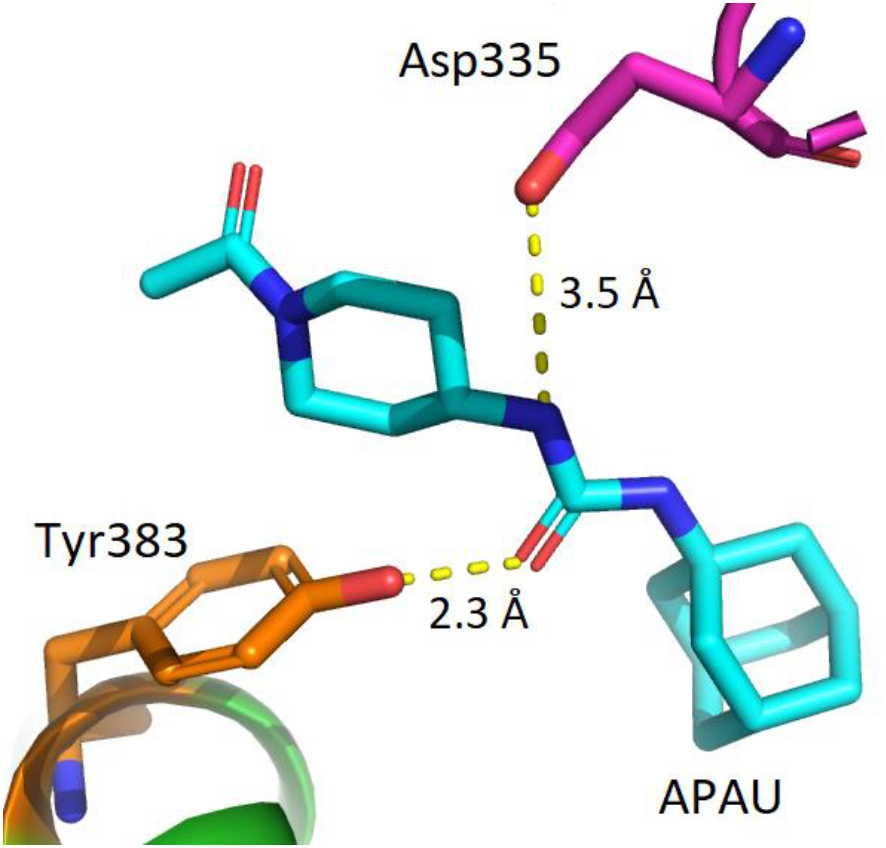
Hydrogen bond formation between APAU and the residues Asp335 and Tyr383 within the active site of sEH.

In this study, the pharmacophore model discussed and the pre-existing structure of APAU are used to create two new sEH inhibitor drug candidates with better docking scores than that of APAU. The calculated absorption, distribution, metabolism, excretion, and toxicity (ADMET) properties of the novel inhibitors are also compared to the ADMET properties of APAU, predicting an improvement as they adhere to Lipinski’s rule of five.^4^ Therefore, these compounds could be potential orally bioavailable drug candidates for various inflammation-related diseases associated with sEH.

## METHODS

The crystal structure of human soluble epoxide hydrolase complexed with TPPU (PDB ID: 4OD0) was obtained through the RCSB Protein Data Bank website.^13^ MakeReceptor under the OpenEye software collection was used to define the area in which the drug potentially binds according to a previously published report.^13,14^ vBROOD 2.0 was used to design a drug candidate via bioisosteric replacement, or the replacement of specified functional groups within the original structure of APAU.^15^ Another drug candidate was created based on biophysical analysis of the pharmacophore and original structure of APAU, with a goal of designing a molecule with an increased number of favorable molecular interactions with the target protein. The two drug candidates were confirmed to not have been previously studied via SciFinder.^16^

Gaussview was used to build the three-dimensional structures of APAU and the two drug candidates and Gaussian 09 was used to optimize these structures.^17^ OpenEye OMEGA2 was used to generate a conformer library containing the drug candidates along with the original structure of APAU.^18^ The conformer library was then docked into the previously designated sEH active site using FRED.^19^

OpenEye FILTER was used to compute the ADMET properties of the three molecules, including Lipinski’s hydrogen bond donors (LHBD), Lipinski’s hydrogen bond acceptors (LHBA), molecular weight (MW), solubility (XLogP), and toxicophores.^20^ PyMOL was used to visualize the protein-ligand interactions and to measure the distances for the interactions between the drug and residues within the active site.^21^ A BLAST search was conducted to find an sEH homolog within a variety of organisms such as beneficial microbes, pathogenic microbes, mice, and various primates.^22^ Jalview was then used to examine the sequences of the homologous proteins to identify any potential off-target effects or to see if the drugs could be tested in preclinical animal studies.^23^ The process of defining the binding site was repeated with the sEH homolog and the candidates were visualized in PyMOL.

## RESULTS AND DISCUSSION

### Analysis of known sEH inhibitors

The structures of the known sEH inhibitors, TPPU, AUDA, and APAU were built and optimized in Gaussian and OpenEye. These structures were analyzed for ADMET properties in Table 1. AUDA and APAU were found to contain one toxicophore due to the presence of the urea pharmacophore. TPPU, containing the same urea group, had an additional three toxicophores as a result of the three alkyl halides in the molecule. Despite the presence of these toxicophores, none of the molecules were reported to form toxic metabolites in any of the studies.^4,5,6^ None of the molecules were found to be aggregators and only AUDA was found to violate Lipinski’s rule of five due to its logP being over five.

**Table 1.** ADMET properties and docking scores for the known sEH inhibitors TPPU, AUDA, and APAU.

| Name | MW (g/mol) | L HBD | L HBA | XLogP | Docking Score |
| --- | --- | --- | --- | --- | --- |
| TPPU | 359.3 | 2 | 6 | 2.82 | -7.96 |
| AUDA | 391.5 | 2 | 5 | 5.93 | -7.75 |
| APAU | 319.4 | 2 | 5 | 1.87 | -6.98 |

The docking scores for the four known sEH inhibitors were computed and are also reflected in Table 1. All three of the sEH inhibitors tested had similar, within one unit, predicted docking scores. APAU and AUDA were found to be more soluble compared to TPPU according to the FILTER report. Due to its high solubility, and lack of Lipinski rule violations, APAU was selected to be used as the base-structure for designing a drug candidate with improved binding affinity. Literature suggests that the residues Asp335 and Tyr383 form tight hydrogen bonds with the urea structure within APAU.^11^ Indeed, in the docked structure, the bond lengths were 2.3 Å between Tyr383 and the carbonyl on the urea group and 3.5 Å between Asp335 and the hydrogen connected to the urea structure. These interactions between APAU and the active site give insight into what should be retained in the structure of newly designed inhibitors.

### Hypothesis driven design of Candidate 1

Previous studies have proved the significance of the primary pharmacophore by showing how the sEH inhibitors bind within the active site.^10^ For this reason, the urea structure was retained in the design of *Candidate 1*. Furthermore, the proposed pharmacophore model in Figure 2 was used as a template for constructing the new drug candidate. When studying multiple sEH inhibitors, it was discovered that drugs containing a trifluoromethoxy phenyl group had more favorable ADMET properties such as better absorption and a longer half-life as a result of slower metabolism and excretion.^25^

As such, it was hypothesized that linking a trifluoromethyl ether to the existing structure of APAU via a non-polar linker such as a three-carbon chain would increase binding affinity and result in more favorable ADMET properties. This new structure, seen in Figure 4, retains the important structural elements of APAU such as the urea group and the piperidine ring. A SciFinder search indicated that no identical structures have been previously studied.

**Figure 4.**
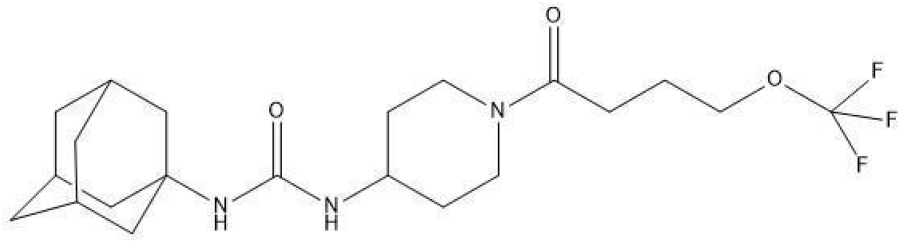
Two-dimensional structure of Candidate 1.

*Candidate 1* had a docking score of −12.08 (Table 2), which is a 73% improvement compared to that of APAU (−6.98). It was hypothesized that the new drug candidate would have similar interactions to those between APAU and the binding site. However, when examining the protein-ligand interactions for *Candidate 1* (Figure 5), the urea group did not bind to Asp335, nor Tyr383 as expected. Instead, the two residues formed hydrogen bonds with the carboxyl group attached to the piperidine (bond length 3.0 Å) and the fluorine on the trifluoromethyl ether (bond length 3.5 Å). The change in docking position is predicted to result in a stronger hydrogen bond between *Candidate 1* and Asp335 as the bond length changed from 3.5 Å to 3.0 Å. Additionally, the different orientations of the drug within the active site allowed for the formation of an additional hydrogen bond. The new hydrogen bond between Tyr466 and the oxygen in the ether group had a bond length of 3.0 Å. This additional interaction and the stronger hydrogen bonds likely contribute to the better predicted binding affinity of *Candidate 1* compared to the original structure of APAU.

**Table 2.** ADMET properties and docking score for *Candidates 1* and *2*.

| Name | MW (g/mol) | L HBD | L HBA | XLogP | Docking Score |
| --- | --- | --- | --- | --- | --- |
| C1 | 431.4 | 2 | 6 | 3.61 | -12.08 |
| C2 | 360.5 | 3 | 5 | 2.64 | -9.76 |

**Figure 5.**
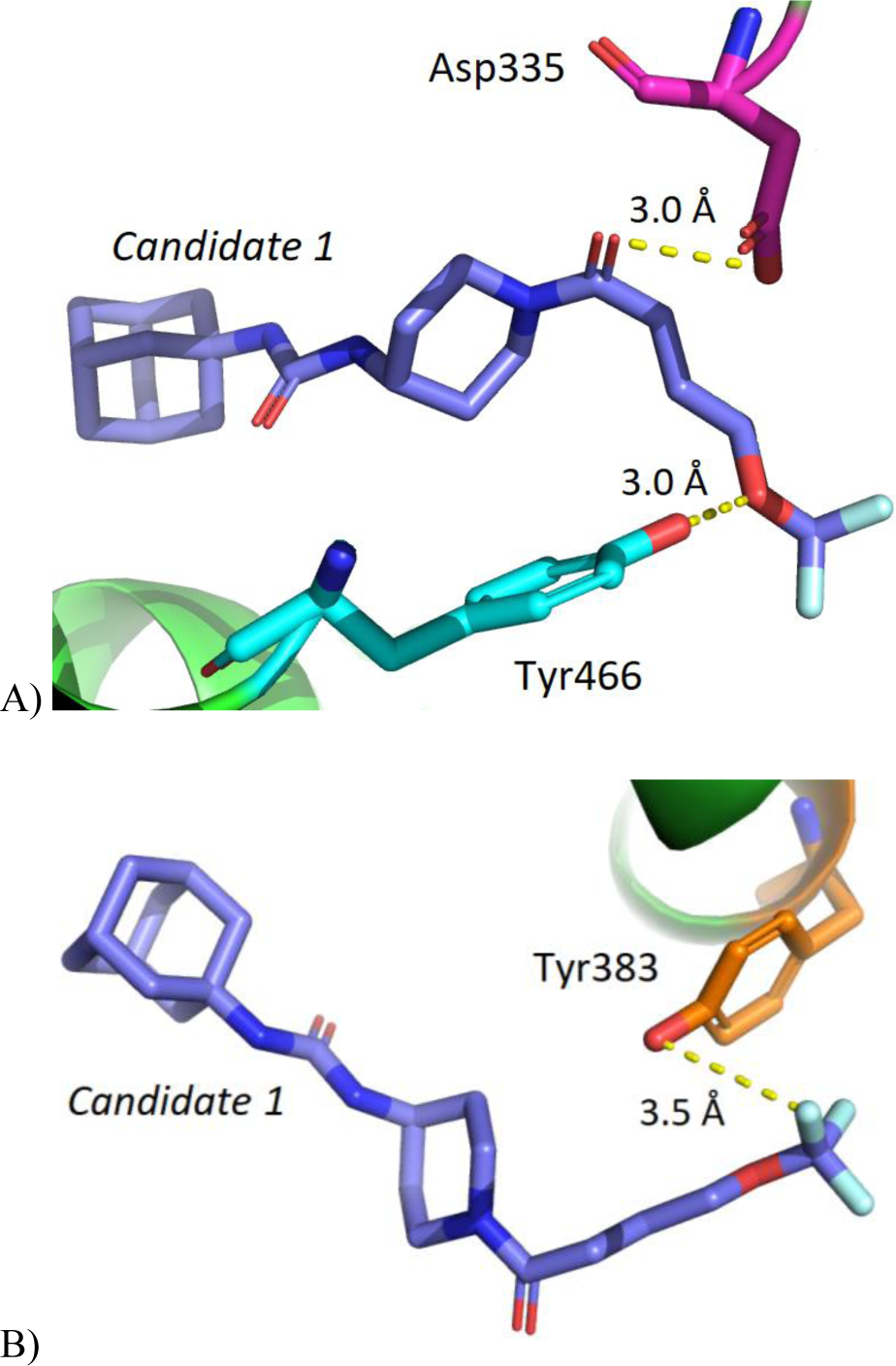
Interactions between *Candidate 1* and the sEH active site. A) Hydrogen bonds between *Candidate 1* and active site residues Asp335 and Tyr466. B) Hydrogen bond between *Candidate 1* and Tyr383.

To determine whether *Candidate 1* will be a good drug candidate, the ADMET properties generated by FILTER (Table 2) were evaluated. *Candidate 1* passed the criteria for Lipinski’s rule of 5, indicating that the drug will be orally bioavailable. Furthermore, the drug had no additional toxicophores compared to TPPU, which also contains a trifluoromethoxy phenyl group similar to the trifluoromethyl ether group in *Candidate 1*. Lastly, *Candidate 1* passed both the aggregator and filter tests, indicating that the drug will not potentially inhibit or activate other proteins within the body in a non-specific manner.^25^ Overall, analysis of *Candidate 1* showed that it is a promising drug candidate with the potential to be a better inhibitor than APAU.

### Computationally driven design of Candidate 2 Candidate 2

was generated using vBROOD 2.0 to find a bioisostere of APAU. To comply with the pharmacophore model in Figure 2, a majority of the original structure of APAU was retained, including the adamantine group, urea group, and the piperidine structure. For these reasons, the acetyl group was chosen for modification.

Multiple analogs were generated and docked into the sEH active site. Of the analogs generated, *Candidate 2* (Figure 6) had the highest binding affinity with a docking score of −9.76, a 40% improvement compared to APAU. In *Candidate 2*, the acetyl group was replaced with a cyclopropane attached to an acetyl group. The cyclopropane addition acted as the non-polar linker, while the acetyl acted as the polar group. These additions adhered to the pharmacophore model proposed in Figure 2. Additionally, the nitrogen in the piperidine ring was protonated. A search on SciFinder indicated that no identical structures have been previously studied. The docking score and ADMET properties calculated using FILTER are reported in Table 2.

**Figure 6.**
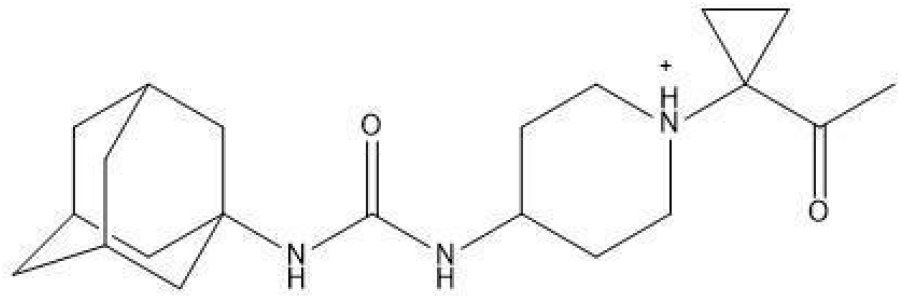
Two-dimensional structure of *Candidate 2*.

Figure 7 shows the interactions between the sEH active site and *Candidate 2*. The bonds between Asp335 and *Candidate 2* have bond lengths of 3.2 Å and 3.0 Å respectively. The hydrogen bond between Tyr383 and the drug has a length of 2.4 Å, and that between His523 and the drug has a length of 1.6 Å. The higher docking score may be the result of an increase in the number of hydrogen bonds being made between Asp335 and the drug candidate and shorter hydrogen bond lengths (3.2 Å and 3.0 Å compared to the original 3.5 Å). Additionally, *Candidate 2* forms a hydrogen bond with His523, which was not observed with the binding of APAU to the active site.

**Figure 7.**
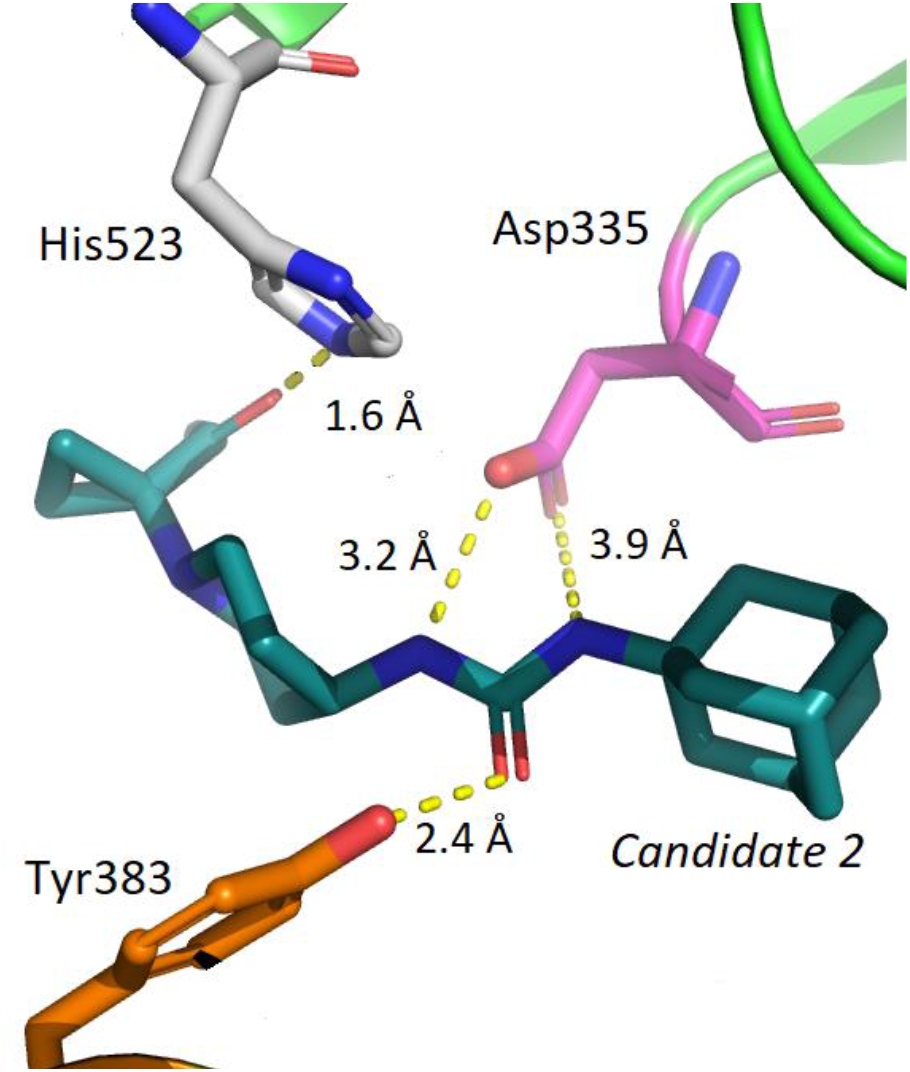
Hydrogen bonds between *Candidate 2* and active site residues Asp335, Tyr383, and His523.

Table 2 reflects the ADMET properties for *Candidate 2*, generated by FILTER. The drug passed the aggregator and filter tests, as well as the criteria for Lipinski’s rule of 5. Similar to *Candidate 1*, there were no additional toxicophores compared to the original structure of APAU, indicating that it is not likely to form toxic metabolites in the body. Analysis of *Candidate 2* showed that the drug is a promising candidate with the potential to have better inhibitory properties than APAU.

### Homolog Analysis

To find out if there were beneficial or harmful off-target effects, a protein alignment search was conducted with the following beneficial and pathogenic microbes: *lactobacillus, Bifidobacterium, Akkermansia muciniphila, Staphylococcus aureus, Haemophilus influenzae*, and *Candida albican* with BLAST. The results showed that all the organisms examined had proteins with less than 30% identity, indicating that the protein sequences in these organisms were not similar to human sEH. The low homology among proteins within the microbes examined indicated that the drug candidates are not likely to bind as there are no comparable binding sites. A secondary BLAST search was conducted to determine which species would be appropriate for testing in preclinical animal models. The search identified homologs with identities above 98% for a large group of nonhuman primates (including, but not limited to *Pan troglodytes, Pan paniscus, Gorilla gorilla gorilla, Pongo abelii, Hylobates moloch, Nomascus leucogenys, Piliocolobus tephrosceles*) and 73.2% identity for the species *Mus musculus*, or the common house mouse. Sequence alignment showed that among all the species listed, multiple residues within the active site of the enzyme were conserved at 99% identity, including the functionally important Asp335 and Tyr383.

The crystal structure for sEH in *Mus musculus* was available (PDB ID: 1CR6) and was used to examine the interactions between the homologous protein and the two drug candidates (Figure 8).^26^

**Figure 8.**
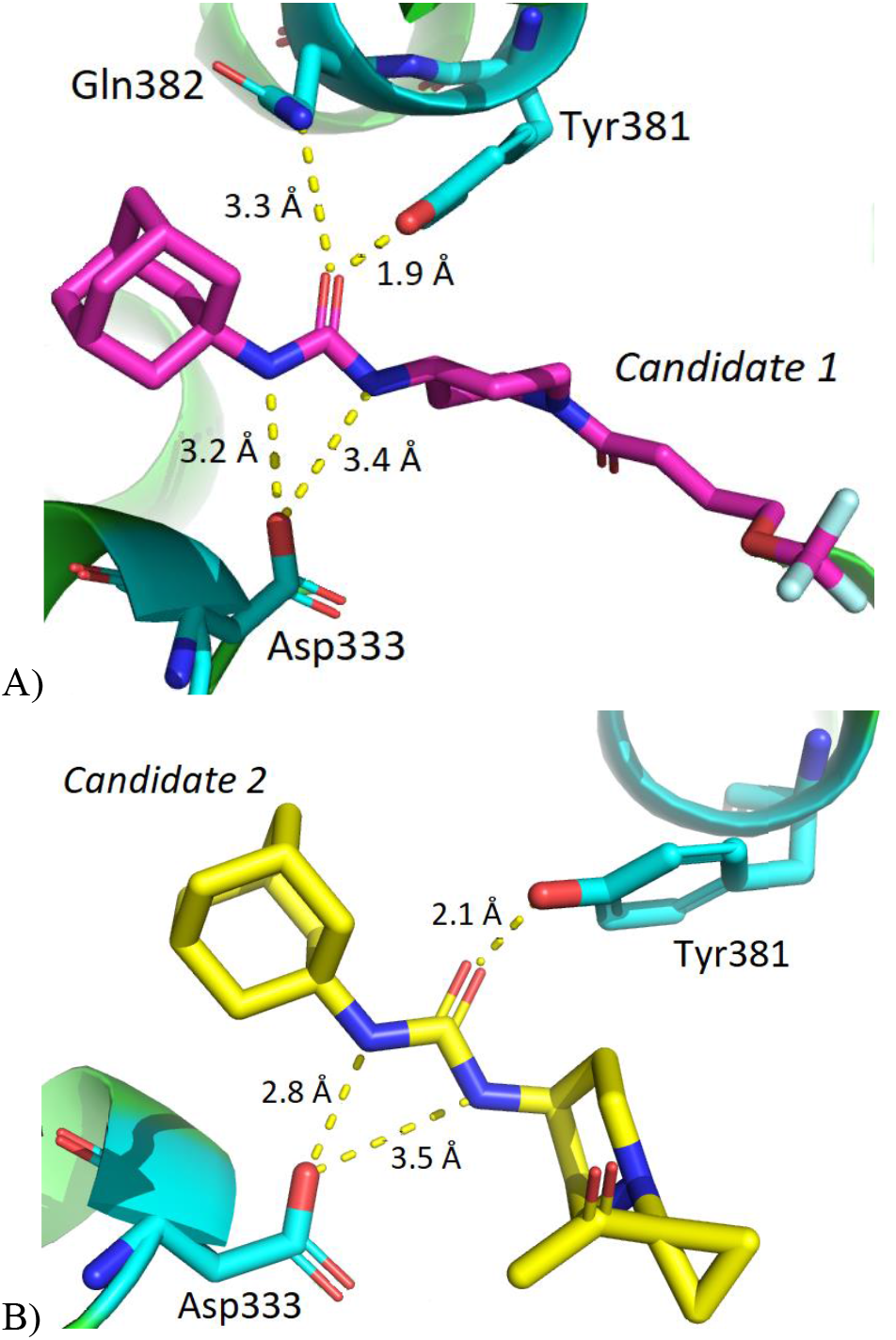
Interactions between the drug candidates and the sEH homolog in *Mus musculus*. A) *Candidate 1* hydrogen bonding with the residues Tyr381, Asp333, and Gln382. B) *Candidate 2* hydrogen bonding with the residues Tyr381, and Asp333.

The docking scores for Candidates 1 and 2 in *Mus musculus* sEH were calculated to be −17.94 and −14.21, respectively. The docking scores for the drug candidates within *Mus musculus* indicate a greater binding affinity compared to the docking scores of the drug candidates within human sEH (−12.08 and −9.76 for Candidates 1 and 2 respectively).

The residues Asp333 and Tyr381 in *Mus musculus* corresponded to the active site residues Asp335 and Tyr383 in human sEH and formed similar hydrogen bonds with the urea structure in the drugs. For *Candidate 1*, two hydrogen bonds were formed with the residue Asp333 with lengths of 3.4 Å and 3.2 Å. *Candidate 2* similarly formed two hydrogen bonds with Asp333, with bond lengths of 2.8 Å and 3.5 Å. Both *Candidates 1* and *2* formed hydrogen bonds with Tyr381, with bond lengths of 1.9 Å and 2.1 Å, respectively. *Candidate 1* was observed to form an additional hydrogen bond with the residue Gln382 with a bond length of 3.3 Å. The Gln382 residue in *Mus musculus* corresponded to the Gln384 residue in human sEH.

The drug candidates interact with residues within *Mus musculus* sEH that correspond to residues in human sEH. The similarity in binding indicates that the sEH in mice has an active site similar to human sEH. The presence of hydrogen bonds between the drug candidates and the similar residues within the murine epoxide hydrolase indicate that the drugs are likely to bind within mice models. Therefore, in preclinical trials for these drug candidates, mice would be appropriate test subjects.

## CONCLUSION

Soluble epoxide hydrolase inhibitors have a wide range of applications as they are currently being studied for a multitude of different diseases. Although none have been approved for human use, it is important to explore a variety of different drug candidates in order to find one with the best properties, specifically pertaining to ADMET properties. The two novel drug candidates designed in this study through both chemically and computationally-driven methods show potential to be potent sEH inhibitors through desirable ADMET properties and by displaying improved docking scores compared to the pre-existing sEH inhibitors.

Protein sequence alignment showed a high similarity in the sEH proteins between species, as the proteins were found to be over 95% similar. Therefore, mouse and nonhuman primate models are appropriate for preclinical pharmacokinetics and pharmacodynamics studies. Testing these drugs in preclinical trials would examine the safety and efficacy of the drugs prior to testing in human clinical trials.

## AUTHOR INFORMATION

### Author Contributions

Research was designed by all authors; all experiments were carried out by C.C. The manuscript was written through contributions of all authors. All authors have given approval to the final version of the manuscript.

## ACKNOWLEDGMENT

Research reported in this publication was supported by UC Davis, the National Science Foundation Award Numbers 1827246, 1805510, 1627539, the National Institute of Environmental Health Sciences of the National Institutes of Health (NIH) under Award Number P42ES004699, UC Davis, NIH Award Number R01 GM 076324-11 and the Rosetta Commons. The content is solely the responsibility of the authors and does not necessarily represent the views of the National Institutes of Health or National Science Foundation. This study was derived from a course based undergraduate research study conducted in Chemistry 130B at UC Davis.

